# Bifidobacterium- and Escherichia-dominant ecological guilds shape distinct microbial metabolism in the gut microbiome of tuberculosis patients

**DOI:** 10.1101/2025.11.24.690090

**Authors:** Milyausha Yunusbaeva, Dina Sabirova, Liliya Borodina, Karina Khasanova, Aliya Yunusbayeva, Radick Altinbaev, Ildar Ganiev, Bayazit Yunusbayev

## Abstract

**Introduction:** Gut microbial community plays a key role in maintaining host immune homeostasis. However, current analytical approaches treat microbiomes at taxon-level and miss community-level functions in the so-called ecological guilds. Therefore, to understand community-level impact on the host, it is essential to reconstruct co-occurring ecological partners and understand their joint metabolic function.

**Methods:** In this study, we first decomposed the fecal microbiota of tuberculosis patients and healthy controls into five key bacterial enterosignatures (ESs) using the non-negative matrix factorization (NMF) approach. This approach allowed us to group individual microbiome members into ecological units that better capture community-level function and correlate with host biomarkers and disease.

**Results and Discussion:** We show that ecological guilds, the so-called enterosignatures, reproducibly describe gut microbiome and provide solid basis for comparison across studies. Namely, our healthy participants had the same mixture of ecological guilds observed among worldwide collections of human donors. Compared to this healthy ecological profile, tuberculosis patients demonstrated unusual enrichment of two bacterial enterosignatures, designated ES-Bifi and ES-Esch that often appear in donors with disturbed or transient gut communities. We inferred metabolic pathways enriched in these two enterosignatures and delineated metabolic processes dominating TB patients’ gut communities. Specifically, we found that ES-Bifi and ES-Esch members use rapid growth strategies based on anaerobic glycolysis that lead to excessive fermentation into acetate and lactate. Higher number of bacteria using anaerobic glycolysis hinted at higher glucose availability. We hypothesize that higher fermentation into lactate may cause acidification and suppressed normal flora evident in patients.

## 1 Introduction

Tuberculosis (TB) remains the deadliest infectious disease, imposing a significant global burden, especially in developing countries. In 2023, 8.2 million people were newly diagnosed with tuberculosis worldwide, an increase from 7.5 million in 2022 and 7.1 million in 2019 (World Health Organization, 2024). One contributing factor to this rise is the variability in treatment response, underscoring the urgent need to understand the host factors that influence differential immune resistance.

The gut microbiota plays a crucial role in maintaining host immune homeostasis and mediates colonization resistance in both the intestine and other distal organs (Budden et al., 2017; Campbell et al., 2023; Libertucci and Young, 2019; X. V. Li et al., 2019). Murine models of tuberculosis have shown that alterations in the gut microbiota can increase host susceptibility to *Mycobacterium tuberculosis* (MTB) infection (Khan et al., 2016). Furthermore, changes in the gut microbiota impair isoniazid-induced MTB clearance by reducing innate immunity and attenuating the CD4+ T cell response to MTB (Khan et al., 2016). Numerous studies investigating the gut microbiome in TB patients have reported shifts in the taxonomic composition of the gut community and dysbiosis (Hu et al., 2019; W. Li et al., 2019, 2024; Luo et al., 2017; S. Wang et al., 2022; Ye et al., 2022; Yu et al., 2023). However, taxonomic changes and disease-associated species cannot tell us much about the impact of the gut microbiome on the host. Without understanding bacterial gene content and metabolic potential, it is difficult to assess the potential impact on the host.

Ecological and microbial interactions are another layer of complexity. In the real gut community, bacteria coexist with their ecological partners, forming so-called ecological guilds. Therefore, to understand their functional impact on the host, it is essential to reconstruct co-occurring ecological partners and understand their joint metabolic potential, which is often overlooked by standard statistical methods. Co-occurring ecological partners, or ecological guilds, can be reconstructed as enterosignatures—a new concept—that, unlike discrete enterotypes (Arumugam et al., 2011), better represent human gut communities. Recently, Frioux et al. (Frioux et al., 2023) demonstrated that the human gut microbiome comprises at least five complementary enterosignatures (ESs), each of which is a stable ecological entity that can be reproducibly recovered from diverse human populations. ESs represent both typical healthy gut microbiomes and atypical microbiomes associated with adverse host health conditions and/or the presence of pathobionts. In this study, we utilized pre-computed enterosignatures representing the global diversity of 5,230 fecal metagenomes from 13 countries and various age groups, referred to as the Gut Microbiome Reference Model (GMR), to decompose the fecal microbiota of TB patients and healthy donors into bacterial enterosignatures. This approach enabled us to identify two key bacterial guilds, designated as ES-Bifi and ES-Esch, that were unusually increased in the gut microbiome of TB patients. The observed increase in these two ESs suggested that the intestines of TB patients have distinct ecological conditions compared to those of healthy individuals. We reconstructed metabolic pathways enriched in these two enterosignatures and predicted the metabolic processes that dominate the gut microbiomes of patients. Finally, we compared our findings with conventional approaches commonly used in gut microbiome research. We demonstrate that modeling microbiomes as ecological guilds enables the detection of microbial processes that are overlooked by traditional techniques and that can have a direct impact on the disease.

## 2 Methods

### 2.1 Study design, samples, and quality control

In this case-control study, we collected 80 freshly frozen fecal samples from 47 healthy adult donors and 33 treatment-naive patients with active pulmonary tuberculosis admitted to the Republican Clinical TB Dispensary in Ufa, Russia, between 2019 and 2021. The metagenomic sequences from 47 healthy controls and a subset of TB patients were previously published in another study conducted by our team (Yunusbaeva et al., 2024). All patients had newly diagnosed active pulmonary TB as confirmed by chest radiography and a positive sputum smear or were positive for *M. tuberculosis* based on the GeneXpert MTB/RIF test without evidence of rifampin resistance. Exclusion criteria included severe comorbidities, hemoptysis, hypoxia, extrapulmonary tuberculosis, history of tuberculosis treatment, use of anti-TB drugs within the past 30 days, pregnancy or breastfeeding, and HIV infection. Fecal samples were collected from TB patients before treatment commenced. Control participants (n = 47) were recruited from staff members working at the Institute of Biochemistry and Genetics of the Russian Academy of Sciences (Ufa, Russia), and were matched for age and sex with the cases. These control subjects had no unexplained symptoms or other conditions, including infectious, autoimmune, or gastrointestinal diseases. All subjects showed no abnormalities on radiographic imaging. Before entering the study, all participants provided written consent.

Fresh stool samples from the participants were collected using sterile containers and immediately stored at -80 °C for further analysis. Bulk DNA was extracted from each donor’s fecal samples using the QIAamp DNA Stool Mini Kit (Qiagen, Netherlands). Sequencing libraries were prepared using the TruSeq DNA PCR-Free Sample Preparation Kit (Illumina, USA), following the manufacturer’s instructions. Deep metagenomic sequencing was performed on the NovaSeq 6000 platform using 150 bp paired-end reads, generating approximately 30 million sequences per donor.

### 2.2 Raw sequence QC and preprocessing

We assessed sequence quality before and after quality trimming and decontamination using FastQC v0.11.9 (Wingett and Andrews 2018). Trimmomatic v0.33 (Bolger et al., 2014) was employed to remove low-quality bases (Q20), and fastp v0.23.2 was used to clip Illumina adapters (Chen et al., 2018). Additionally, tandem repeats were removed using Tandem Repeats Finder v4.09 (Benson, 1999). Finally, the KneadData v0.10.0 pipeline from the bioBakery toolset was utilized to decontaminate reads originating from the human genome, transcriptome, and microbial RNA (McIver et al., 2018). Specifically, we used the human reference genome (build hg37), the human reference transcriptome (build hg38), and the SILVA ribosomal RNA reference databases.

### 2.3 Enterosignature analysis of gut microbiome composition

To identify biologically meaningful patterns in gut microbiome composition, we applied enterosignature analysis using the online enterosignature tool (https://enterosignatures.quadram.ac.uk/) based on the r207 GTDB database. Enterosignatures represent latent components that capture co-abundance patterns of microbial taxa across samples, providing a dimension-reduced representation of microbiome community structure. The tool decomposed the microbiome composition into five distinct enterosignatures (ES-Bact, ES-Bifi, ES-Esch, ES-Prev, and ES-Firm), each representing a specific microbial community configuration. For each sample, we calculated the relative contribution of each enterosignature, with the sum of all signatures normalized to 1.0 per sample. Sample quality was assessed using cosine similarity between the original species abundance profile and its reconstruction from the enterosignature decomposition. We classified samples based on model fit quality and identified the dominant enterosignature for each sample as the signature with the highest relative abundance. We performed comprehensive analyses to characterize enterosignature patterns across treatment-naive TB patients and healthy controls. Distribution patterns were visualized using stacked bar plots, with samples ordered by clinical group, dominant enterosignature, and clustering patterns. Hierarchical clustering using Euclidean distance with Ward’s linkage method was applied to identify natural groupings of samples within the TB patient group based on their enterosignature profiles.

Statistical differences in enterosignature abundances between treatment-naive TB patients and healthy controls were tested using Wilcoxon rank-sum tests, applying Benjamini-Hochberg correction for multiple comparisons. Permutational multivariate analysis of variance (PERMANOVA) based on Aitchison distance was used to test overall compositional differences between groups. We additionally analyzed samples with high abundance of specific enterosignatures, particularly focusing on ES-Esch and ES-Bifi, using a threshold of 3% relative abundance to identify samples dominated by these signatures.

### 2.4 Bacterial community diversity and principal component analysis

Shannon alpha and beta diversity measures were estimated using the microbiome R package (Lahti and Shetty, 2017). Differences in Shannon alpha diversity between patients and controls were tested using the two-sample Kolmogorov-Smirnov test implemented in the ks.test() function of the stats R package (R: A Language and Environment for Statistical Computing, 2022). To visualize gut community differences between donors using major axes of microbial variation, we applied principal component analysis (PCA) on the species abundance matrix. To adapt sparse data on relative abundance for principal component analysis, we imputed zeros in the relative abundance table using the cmultRepl() function in the zCompositions R package version 1.4.0-1 (Palarea-Albaladejo and Martín-Fernández, 2015). Next, to take compositionality into account, the relative abundance table was transformed using the center log ratio (CLR) prior to PC analysis. PC analysis was carried out using the prcomp() function in the stats R package (R: A Language and Environment for Statistical Computing, 2022).

### 2.5 Inference of taxonomic content of the gut microbiome and between-group comparisons

Before downstream analyses, all bacterial taxa observed only in a single donor and at a relative abundance of less than 0.0001 were discarded. The choice of an abundance threshold ensured that we kept rare bacterial species that can impact human disease, as demonstrated in recent work (Chriswell et al., 2022). Metaphlan v4.0 tool was used to infer both known bacterial species and strains, as well as uncharacterized bacterial species. Uncharacterized bacterial species in the Metaphlan v4.0 tool are defined in terms of species-level genome bins (SGBs) using so-called metagenomically assembled genomes (MAGs) (Blanco-Míguez et al., 2023). To facilitate comparison with published data, we analyzed the relative abundances of bacterial taxa in sampled groups (TB patients and controls) at different taxonomic levels by aggregating strains and species into genera, families, orders, classes, and phyla. We used linear discriminant analysis with effect size estimation (LEfSe), implemented in the microbial R package, to identify bacterial taxa that characterize the differences between patients and controls. We also calculated the average proportion of each taxon for patients and controls at different taxonomic levels (e.g., phylum and class). A statistical difference in proportions between groups was tested using the ANCOM statistical framework (Mandal et al., 2015), employing the false discovery rate (FDR) approach to adjust p-values. When we tested average proportions using ANCOM, we were interested in comparing dominant taxa between groups. Therefore, we required bacterial taxa to be present in at least 25% of donors at an abundance of 5% or greater.

### 2.6 Inference of the microbial metabolic capacity of the gut microbiome and between-group comparisons

The metabolic capacity of the microbiome, often referred to as the metabolic profile (potential), was inferred using the HUMAnN 3.6 framework using default settings. Briefly, HUMAnN 3.6 compares the sequences from donor metagenomes to known microbial genomes and genes to infer known metabolic reactions and aggregate them into known microbial pathways.

Specifically, for bioinformatic queries, HUMAnN 3.6 utilizes the ChocoPhlAn 3 database, which comprises 99,200 annotated reference genomes from 16,800 microbial species in the UniProt database (as of January 2019) (UniProt Consortium, 2019) and the corresponding functionally annotated 87.3 million UniRef90 gene families (Suzek et al., 2015). For each input donor (fecal) metagenome, HUMAnN 3.6 provides bacterial-specific genes and pathway abundance whenever genes can be attributed to known bacterial species in the database. Gene families are annotated by default using UniRef90 definitions, and pathways are annotated using MetaCyc definitions (Caspi et al., 2020). Whenever inferred genes cannot be attributed to a specific bacterial species, gene and pathway abundances are reported without specifying their bacterial provenance. To identify metabolic pathways that are differentially abundant in patients versus controls, we used linear discriminant analysis with effect size estimation (LEfSe), implemented in the microbial R package. For pathways that showed strong enrichment in patients (those with the largest effect size and the highest LDA score), we also inferred the bacterial species, genera, and higher-level taxa that were major contributors to that pathway.

## 3 Results

### 3.1 Strength of the relationship between the gut microbiome and disease

Earlier studies have described taxonomic alterations in TB patients, but without reporting standardized effect size estimates. Effect size estimates are essential to a) understand the strength of the relationship between disease and the gut microbiome; b) facilitate comparison between studies using standardized measures, and c) perform power analysis. In this study, we used Cohen’s d to assess the effect size of tuberculosis on the key microbiome readouts (Rahman et al., 2023). Namely, measures of community diversity, principal components (PCs) of taxonomic variation, and principal components of metabolic pathway variation observed in the sample. As a rule of thumb, Cohen’s d values of 0.2, 0.5, and 0.8 are considered as “small,” “medium,” and “large” effect sizes, respectively (Rahman et al., 2023).

Our estimates showed that tuberculosis have an exceptionally strong (Cohen’s d = 3.855) effect size on taxonomic variation (Supplementary Figure 1), which is much stronger than for chronic inflammatory diseases (Autoimmune diseases, Cohen’s d ∼ 0.2) and fungal overgrowth (Cohen’s d ∼ 0.25), by far the strongest factor impacting gut microbiome (Rahman et al., 2023). Accordingly, the strong impact on the species content shaped the top axis of taxonomic variation, PC1, which clearly separated patients from controls in the PCA plot (Supplementary Figure 2). It should be noted that observed taxonomic alterations in TB patients were accompanied by a moderate effect on taxonomic diversity, as indicated by the Shannon index (Cohen’s d = 0.368).

Next, we found that tuberculosis had a notably strong effect on the functional capacity of the gut microbiome, specifically in terms of variation in microbial metabolic pathways, as summarized by PC1 and PC2 (Cohen’s d = 0.841 and Cohen’s d = 2.009, respectively) (Supplementary Figure 1). These two top principal components (PCs) of metabolic pathway variation jointly separated TB patients from controls in PC analysis (Supplementary Figure 2). Unlike taxonomic diversity, diversity of metabolic pathways in the gut microbiome was impacted much strongly (Cohen’s d = 1.044) by tuberculosis status (Supplementary Figure 1). In summary, our study found that tuberculosis had a significant impact on the gut microbiome in the sample, and our moderate sample size (N=80) provided sufficient statistical power (over 80%) to detect biologically meaningful differences in the species and metabolic pathway content of the gut microbiome (Supplementary Figure 1).

While tuberculosis outcompeted many of the known factors influencing the gut microbiome, we took measures to minimize variability of age and sex between patients and controls. We compared the demographic characteristics of patients with those of healthy donors before their inclusion in the study cohort. Supplementary Table 1 presents the baseline characteristics of all participants. We found no significant differences in sex or age between the TB patients and controls (p < 0.05, Supplementary Table 1). The median age of the patients was 43 years (IQR 34.5–55), and male TB patients slightly outnumbered females (57.6% vs. 42.4%).

### 3.2 The structure of ecological guilds indicates an altered niche in the patient’s gut

Bacteria live in communities, yet our analytical approaches treat them in isolation, which hinders our ability to decipher their community-level functions. To better understand gut microbial biology and function, we considered gut bacteria in our donors by reconstructing microbial ecological guilds, known as enterosignatures (ESs). Enterosignatures represent highly reproducible bacterial guilds observed in human gut microbiomes across diverse ethnicities worldwide (Frioux et al., 2023). To facilitate cross-study comparison, we used pre-computed enterosignatures representing the global diversity of 5,230 fecal metagenomes from 13 countries and various age groups, referred to as the Gut Microbiome Reference (GMR) (Frioux et al., 2023). Specifically, to identify recurrent enterosignatures in our patients and controls, we applied non-negative matrix factorization (NMF, https://enterosignatures.quadram.ac.uk/) to the relative abundances of genera observed in our donors (Figure 1A). Notably, microbiomes in our samples contained the same five ESs identified earlier in a global sample of donors: ES-Bact (primarily characterized by the genera *Bacteroides* and *Phocaeicola*), ES-Bifi (*Bifidobacterium* and *Streptococcus*), ES-Prev (*Prevotella*), ES-Esch (*Escherichia* and *Citrobacter*), and ES-Firm (genera within the phylum *Firmicutes*). While ESs are designated by the dominating genera, in reality, each enterosignature is composed of a unique combination of genera (Figure 1C). Often, one or two genera dominate ESs and strongly correlate with the ecological guild (for instance, ES-Prev (*Prevotella*)), but some ESs consist of multiple genera with weaker associations (Figure 1C). Members of the same ES, i.e., bacterial guild, co-occur and exhibit consistent behavior since they collaborate to exploit the resources of the same ecological niche.

**Figure 1.**
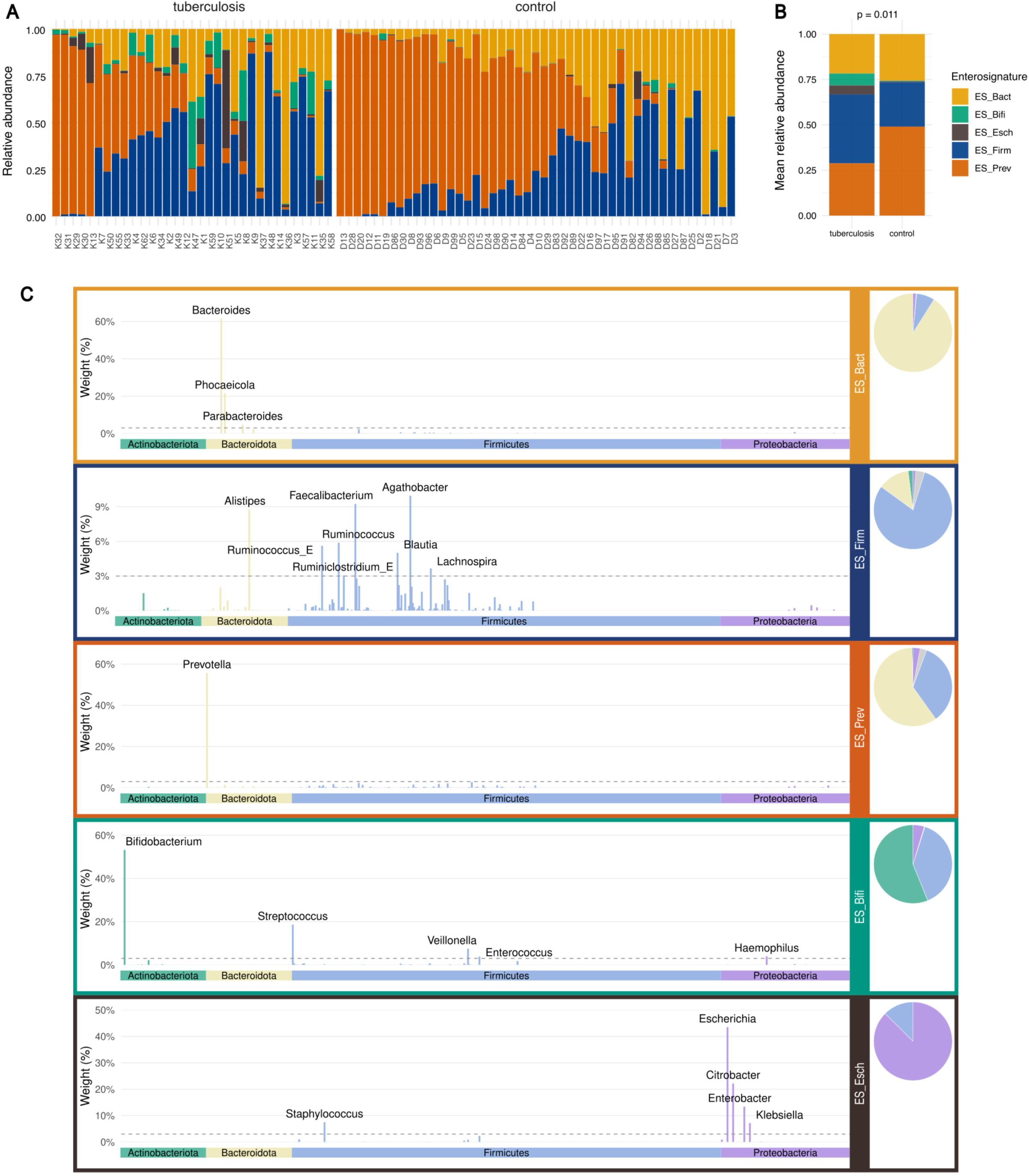
Composition of enterosignatures (ESs) and their diversity in samples. **(A)** The relative abundances of the five ESs in samples from TB patients and controls. **(B)** Average abundances of the five ESs in TB patients and controls. (С) Relative genus-level composition of the five ESs identified as the optimal NMF solution.

As can be seen in Figure 1A, healthy controls in our sample harbour the same combination of enterosignatures (ES-Bact, ES-Prev, and ES-Firm) observed among (“primary ESs”) worldwide collection of healthy adults following a western diet (Frioux et al., 2023). Similar to the worldwide collection, our healthy participants exhibited multiple enterosignatures, which likely correspond to multiple ecological niches within the human gut (Frioux et al., 2023). Quantitatively, 99% of the gut microbes among our healthy controls belonged to the three adult human enterosignatures (ES-Bact, ES-Prev, and ES-Firm) (Figure 1B). Notably, two out of five enterosignatures, ES-Bifi and ES-Esch, were almost absent, or present in a few donors.

When compared to healthy donors, the gut microbiomes of TB patients were more heterogeneous, had shifted proportions of adult-specific enterosignatures, and exhibited a higher occurrence of two ecological guilds that are rare in healthy adults (p = 0.01) (Figure 1A, B). Specifically, TB patients had an elevated proportion of ES-Firm (38% vs. 24.4%, p = 0.04), but a depleted proportion of ES-Prev (28.8% vs. 49%). A distinctive feature of the gut microbiome of TB patients was high occurrence of ES-Bifi (6.6% vs. 0.4%, p = 1.1 × 10^-11) and ES-Esch (4.9% vs. 0.6%, p = 2.1 × 10^-9). In the worldwide collection of gut microbiomes, the ES-Bifi and ES-Esch were more characteristic of early childhood, sick individuals, and the elderly, in contrast to ES-Bact, ES-Prev, and ES-Firm, which were considered “primary ESs” in adult microbiomes (Frioux et al., 2023; Guner et al., 2009). Interestingly, the defining member of the ES-Esch, *Escherichia coli,* has been often observed in TB patients with elevated abundance (Ding et al., 2022; Luo et al., 2017; Séraphin et al., 2023; Yu et al., 2023). Evidence on bifidobacteria, which dominate ES-Bifi, is controversial. Some studies reported decreased abundance (Wang et al., 2022), while others demonstrated increased abundance of *Bifidobacteria* in TB patients (Han et al., 2024). It appears that the gut microbiome in TB patients has two distinctive features - an elevated proportion of ES-Firm, and frequent occurrence of two enterosignatures, ES-Bifi, and ES-Esch, that are rare among adult healthy individuals. According to Frioux et al., gut microbiomes dominated by ES-Firm likely represent a transitional state when gut ecosystems are disturbed by intensive antibiotic therapy. It is hypothesized that ES-Firm is unable to form a stable community alone and often complements other ESs, particularly ES-Bact (Frioux et al., 2023). In agreement with this hypothesis, we never observed monodominance of ES-Firm among our healthy donors. In contrast, some TB patients had a gut community with over 75% ES-Firm, indicating possibly unstable monodominance (Figure 1A). Finally, the two unusual ecological guilds that are frequently found in TB patients (ES-Bifi and ES-Esch) were unevenly distributed and were found exclusively in combination with other ESs. ES-Bifi was present in 75.8% of samples (25 of 33), whereas ES-Esch was detected in 39% of samples (13 of 33).

### 3.3 Conventional taxonomic analysis operates with fragments of ecological guilds

To compare guild-based analysis with conventional taxonomic approaches, we analyzed gut taxonomic content using a popular linear discriminant analysis, LEfSe. In our dataset, LEfSe identified 291 differentially abundant (adjusted p-value ≤ 0.05 and LDA score ≥ 2.5) taxa (species and subclades defined by MAGs), associated with patients or controls (Supplementary Table 2, Supplementary Figure 3). Apparently, individual species have only a marginal association with disease and a small impact on the host. The observed large number of TB-associated species suggests that one has to focus on understanding joint community-level function. Nevertheless, for the sake of comparison, we first emphasized the strongest signals by highlighting bacterial species with an LDA score of 4.5 or higher (Figure 2A). Except for *Escherichia coli and Klebsiella pneumoniae*, other top TB-enriched species, such as *Blautia wexlerae, Prevotella copri clade C,* and *Akkermansia muciniphila,* are frequent gut commensals (Figure 2А). These TB-enriched species, on average, had higher abundances in patients than in controls (Figure 2B); however, it is unclear how to interpret all these changes. To aid interpretation, we reasoned that differentially abundant species represented fragments of ecological guilds. Indeed, when we attribute individual species to enterosignatures (Figure 1C), we found that disease and health-associated taxa were assigned to distinct ecological guilds. Most of the health-associated species represented members of the three primary adult enterosignatures (ES-Bact, ES-Prev, and ES-Firm) (Figure 2A). In contrast, patient-associated species belonged to the ES-Bifi, ES-Esch, and ES-Firm microbial guilds, which are highly unusual for healthy adult microbiomes. When comparing guild-level and taxon-level analyses, two important points are worth mentioning. First, conventional taxon-level analysis essentially highlights fragments of microbial communities (Figure 2A). As a result, members of ecological guilds appear as a random mixture, which obscures their co-occurrence patterns and the guild-level functions they serve. Hence, downstream analyses may overlook community-level functions that are relevant to the host, such as microbial products and metabolic pathways.

**Figure 2.**
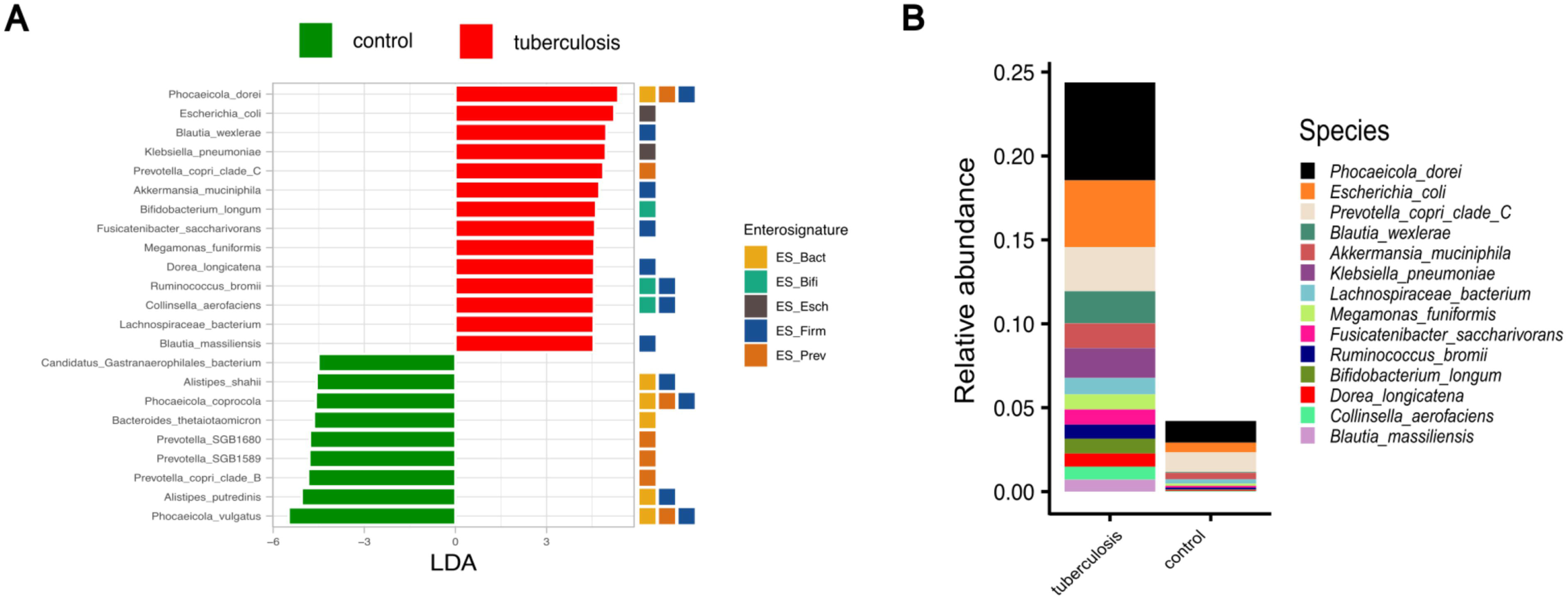
Taxonomic composition of the fecal microbiota of TB patients and healthy controls. **(A)** Barplot depicting top bacterial species (LDA scores > 4.5) discriminating between TB patients and controls. The colored squares to the right of the barplot represent the ESs that contain the specified bacterial species. **(B)** Averaged abundances of the top TB-associated species in TB patients and controls (LDA scores > 4.5). The total height of the stacked bar plot represents the fraction of each species relative to the total taxa present in the gut microbiome of TB patients.

### 3.4 The metabolic profile of the ES-Esch and ES-Bifi enterosignatures in the gut microbiome of TB patients

Since TB patients were unusually enriched in ES-Esch and ES-Bifi, we were interested in delineating metabolic pathways characteristic to these ecological guilds. To do this, we first identified TB patients with an elevated proportion (relative abundance higher than 3%) of these two enterosignatures. Altogether, eight and 17 TB patients had an elevated proportion of ES-Esch and ES-Bifi, respectively. By focusing on these patients, we used gene assemblages in their gut microbiomes to reconstruct metabolic pathways using HUMAnN 3.6 tool (Beghini et al., 2021). Figures 3A and 3B illustrate the metabolic pathways enriched (LDA score ≥ 3) in patients who carry ES-Esch and ES-Bifi. Both patient subgroups exhibited an overabundance of the ANAGLYCOLYSIS-PWY pathway, which facilitates energy generation through anaerobic glycolysis (LDA score > 5) (Figures 3A and 3B). This signal of overabundance does not indicate higher activity of the pathway, but rather an increase in the number of bacteria that carry genes for anaerobic glycolysis. Increased number of bacteria that utilize anaerobic glycolysis may indicate a higher availability of simple sugars in the large intestine of TB patients, enriched with the ES-Esch and ES-Bifi. This finding is notable since healthy digestion must leave only a minimal amount of simple sugars in the large intestine, and homeostatic colon bacteria rely on degrading complex polysaccharides. Therefore, one would expect high abundance of bacterial taxa and their genes (assembled into pathways by HUMAnN 3.6 tool) for degrading complex polysaccharides from three major sources, such as plant-derived glycans, animal-derived glycogen, and host-derived mucin. In this regard, another notable finding is the positive correlation between the ES-Esch enterosignature and elevated blood glucose levels in TB patients compared to controls (Figure 3C). Given our interpretation is valid, the reason for simple sugar availability in the large intestine is unclear. In any case, anaerobic glycolysis is one of the strategies in the gut community that enables rapid cell growth, whereby bacteria quickly convert glucose into energy and metabolic precursors. Our analyses also show that an increased amount of bacterial cells relying on anaerobic glycolysis is paralleled by increased abundance of genes and biosynthetic pathways to synthesize nucleotides and amino acids. For example, pathways to synthesize nucleotides, such as pyrimidine pathways (PWY-5686, PWY-7790, PWY-7199), purine pathways (PWY-6121, PWY-6122), as well as pathways to synthesize amino acids, such as ARGININE-SYN4-PWY, PWY-2942, GLUTORN-PWY, ILEUSYN-PWY, and HISTSYN-PWY (Figures 3A and 3B).

**Figure 3.**
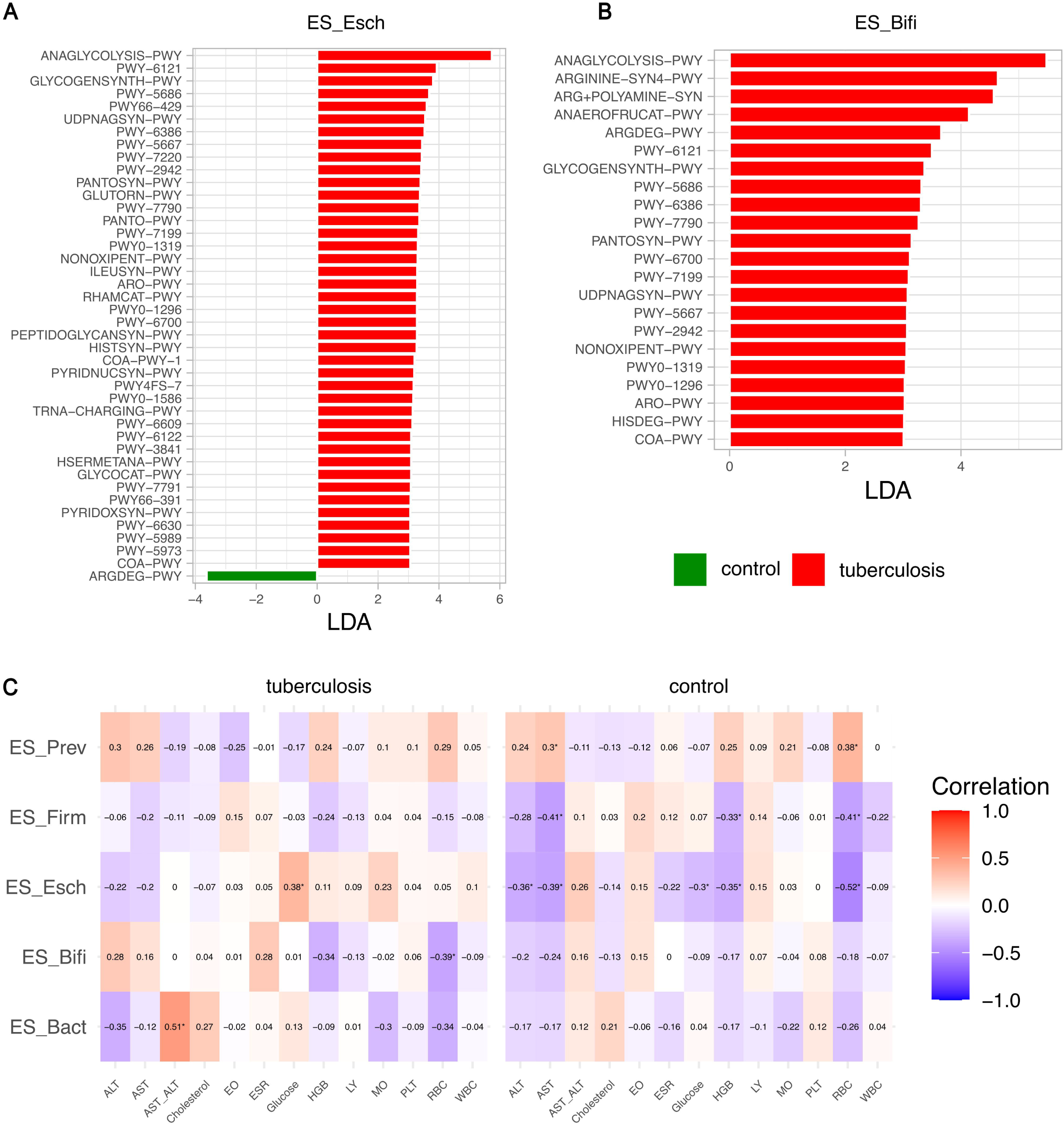
Metabolic composition of the fecal microbiota of TB patients and healthy controls. **(A), (B)** LDA index plots depicting major metabolic pathways (LDA index > 2,5) in fecal samples fortified with ES-Esch **(A)** or ES-Bifi **(B)** from TB patients (red) and healthy controls (green). LDA score plots were generated using the LEfSe analysis. The length of the bar column represents the LDA score. **(C)** Correlation heatmap analysis of blood biochemical indicators and different enterosignatures in TB patients and controls. Red indicates a positive correlation, while blue indicates a negative correlation (P < 0.05). ALT, alanine transaminase; AST, aspartate transaminase; AST_ALT, ratio between the concentrations of two enzymes, aspartate transaminase (AST) and alanine transaminase (ALT); EO, eosinophils; ESR, An erythrocyte sedimentation rate; HGB, hemoglobin; LY, lymphocytes; MO, monocytes; PLT, platelets; RBC, red blood cells; WBC, white blood cells.

Increased number of pathways (number of bacterial gene carriers) supporting cell growth was not surprising and excess bacteria using anaerobic glycolysis hinted at altered flux of simple sugars. In the anaerobic colon, fermentation is the primary mode of energy generation. Therefore, we were interested in fermentation types that are prevalent in order to understand changes in the patient’s colon. We noted that patients with increased ES-Bifi enterosignature show increased gene abundance for homolactic fermentation (Figure 3B). Bacteria that use homolactic fermentation produce (S)-lactate as the sole end product of glucose metabolism. As noted before, increased abundance of pathways means an increased number of species carrying genes for that pathway. Hence, the observed increase in the abundance of homolactic fermentation and anaerobic glycolysis in the ES-Bifi-enriched stool samples may reflect an increased capacity of the intestinal microbiota in these TB patients to acidify the intestinal environment (Wolfe, 2015). It should be noted that ES-Bifi enterosignature contains not only bifidobacteria, which is typically beneficial, but also species from genera such as *Streptococcus, Veillonella, Haemophilus, Enterococcus*. Species from these genera are capable of thriving in acidic conditions and metabolizing lactic acid into short-chain fatty acids, carbon dioxide, and various other metabolites. Essentially, ES-Bifi consists of lactate-tolerant and lactate-producing bacteria. Our correlation analyses between microbial enterosignatures and clinical blood parameters identified a negative association between ES-Bifi and red blood cell counts (Figure 3C). The combination of reduced hemoglobin concentrations, mycobacterial infection, and inflammatory processes leads to tissue hypoxia, subsequently inducing a metabolic shift from aerobic to anaerobic pathways. This metabolic shift leads to the accumulation of lactic acid in the bloodstream, thereby increasing the acidity of both blood and tissues. Thus, our results demonstrate an association between decreased red blood cell counts in TB patients and the presence of ES-Bifi within the gut microbiome.

Finally, we assessed whether grouping bacteria into enterosignatures helps interpretation when compared to conventional approach. We compared metabolic pathways enriched in ES-Esch and ES-Bifi to those enriched in all TB in patients without defining enterosignatures. When gut microbiomes were not classified into enterosignatures, we found 222 metabolic pathways to be differentially enriched either in TB patients or controls (adjusted p-value ≤ 0.05 and LDA score ≥ 2). Of these, 205 pathways were overrepresented in the microbiomes of TB patients (p ≤ 0.05; Supplementary Table 3, Supplementary Figure 4), compared to only 17 pathways enriched in controls. As shown in Supplementary Figure 4, the most enriched pathways in the gut microbiome of TB patients are anaerobic glycolysis (ANAGLYCOLYSIS-PWY), superpathway of arginine and polyamine biosynthesis (ARG+POLYAMINE-SYN), glycogen biosynthesis I (GLYCOGENSYNTH-PWY), pyruvate fermentation to acetate and (S)-lactate I (P41-PWY), pentose phosphate pathway (NONOXIPENT-PWY and PENTOSE-P-PWY), and glycolysis superpathway along with the Entner-Doudoroff pathway (GLYCOLYSIS-E-D) (Supplementary Figure 4, Supplementary Table 3). When we annotated metabolic pathways specific to ES-Esch and ES-Bifi, it became evident that pathways enriched in the TB patients correspond to pathways carried by bacterial species from ES-Esch and ES-Bifi (see black and turquoise squares in Supplementary Figure 4). In fact, most of the metabolic pathways associated with the gut microbiome in TB patients can be explained by increase in bacterial species belonging to ES-Bifi and ES-Esch enterosignatures. Furthermore, we noticed that conventional approach, without defining enterosignatures, failed to detect enrichment of homolactic fermentation that was detected when we defined ES-Bifi enterosignature (Figure 3B). Identification of homolactic fermentation helped us to infer the type of fermentation that was increased in subgroups of patients and hypothesize alterations in their large intestines. In sum, it can be concluded that the metabolic and taxonomic profile of the intestinal microbiota in tuberculosis patients is primarily driven by representatives of ES-Bifi and ES-Esch. The bacterial communities that form these enterosignatures are characterized by active energy production through anaerobic glycolysis, followed by fermentation to acetate and lactate.

## 4 Discussion

In this study, we first decomposed the fecal microbiota of TB patients and healthy controls into bacterial enterosignatures. We utilized precomputed enterosignatures representing the global diversity of 5,230 fecal metagenomes from 13 countries, referred to as the Gut Microbiome Reference Model (GMR). This approach enabled us to group individual microbiome members into ecological functional units, or “enterosignatures,” providing a more ecologically valid method for identifying key patterns between the microbiome and host phenotype. Each enterosignature represents a stable ecological group of closely interconnected and coexisting bacteria that possess molecular mechanisms effectively “link” them together to perform a specific functional role in the gut ecosystem (Frioux et al., 2023). Fecal samples from most participants predominantly contained three enterosignatures: ES-Bact, ES-Prev, and ES-Firm. According to Frioux et al. (2023), ES-Bact and ES-Prev constitute the stable core of a healthy human microbiome, whereas ES-Firm is likely successional and was rarely found in isolation within a sample (Frioux et al., 2023). Although the NMF approach models coexisting enterosignatures that collectively represent a particular fecal microbiome—rather than assigning each sample to a single enterotype—the ES-Bact, ES-Prev, and ES-Firm signatures likely correspond to community types initially described as gut enterotypes (Arumugam et al., 2011). This correspondence indicates a strong biological signal exhibited by these bacterial guilds.

A distinctive feature of fecal samples from TB patients is a statistically significant increase in the abundance of the ES-Bifi, ES-Esch, and ES-Firm. This observed increase suggests that the intestines of TB patients differ ecologically from those of healthy individuals. According to the literature, ES-Esch more accurately characterizes the intestinal microbiome of children, particularly premature infants, formula-fed infants, and children with necrotizing enterocolitis (Guner et al., 2009; Frioux et al., 2023). A high abundance of *E. coli* is also typical in the gut microbiome of older adults and is associated with immunosenescence and diminished host control over the intestinal ecosystem (Fu et al., 2010; Li, Wu, et al., 2025). A high abundance of *E. coli* has been reported in the gut microbiomes of patients with tuberculosis (Yunusbaeva et al., 2024; Yu et al., 2023; Lin et al., 2024), tuberculous meningitis (S. Li et al., 2022), and other diseases (Mirsepasi-Lauridsen et al., 2019; Hoffman et al., 2014). Furthermore, some studies have shown that an increased presence of *E. coli* in the gut microbiome is linked to higher post-meal blood glucose levels and poorer insulin sensitivity (Ju et al., 2023; Chavkin et al., 2021). A study conducted by Madacki-Todorovic and colleagues presented evidence for the direct effect of insulin on increased metabolic activity of *E coli* in association with its biofilm-forming capability (Madacki-Todorovic et al., 2018). Our results also indirectly confirm the correlation between blood glucose levels and the presence of ES-Esch in the gut of TB patients. Additionally, the analysis of metabolic processes characteristic of ES-Esch indirectly supports the availability of glucose in the colon of these patients. Specifically, we observed an increase in the number of species that acquire energy for rapid growth using metabolic pathways that rely on simple sugars.

An intriguing finding of our study is the identification of the ES-Bifi enterosignature in the gut microbiome of TB patients. This result may seem counterintuitive, as *Bifidobacterium* is widely recognized for its health-promoting properties and is commonly used as a probiotic (J. Chen et al., 2021; Hidalgo-Cantabrana et al., 2017; Ku et al., 2024; O’Callaghan and van Sinderen, 2016). However, a large-scale study involving 1,803 Japanese individuals reported that a *Bifidobacterium*-enriched microbiota was significantly associated with a higher prevalence of inflammatory bowel disease, cardiovascular disease, and diabetes (Takagi et al., 2022). Furthermore, a recent study identified specific *Bifidobacteriaceae* taxa as potential biomarkers of disease/health status (Pasolli et al., 2024). A study conducted on patients with pre-extensively drug-resistant tuberculosis also revealed higher levels of *Enterobacteriales, Bifidobacteriales, Verrucomicrobiales*, and *Lactobacillales* within the coliform flora (Shi et al., 2022). Conflicting findings regarding the role of *Bifidobacterium*-rich microbiota in health and disease may be clarified through an enterosignature-based approach. The ES-Bifi is notably enriched not only in bifidobacteria but also in other potentially pathogenic or opportunistic taxa, including *Streptococcus, Veillonella, Haemophilus*, and *Enterococcus*, which may contribute to its association with disease phenotype, potentially modulating or counteracting any beneficial effects of *Bifidobacterium*. Analysis of the ES-Bifi metabolic profile revealed enrichment of pathways associated with energy production from various sources, along with increased homolactic fermentation leading to lactate formation. Lactate, produced through bacterial fermentation in the gastrointestinal tract, plays distinct metabolic and signaling roles in gut dysbiosis and certain metabolic disorders (J. Li, Ma, et al., 2025). Notably, excess D-lactate produced by certain intestinal bacteria can enter the bloodstream and cause metabolic acidosis, particularly in patients with malabsorption syndrome (Levitt and Levitt, 2020; J. Li, Ma, et al., 2025). In the context of tuberculosis, elevated serum lactate levels have long been associated with higher mortality during TB treatment (Ntambwe and Maryet, 2012; Im et al., 2021). Mechanistically, lactate, on the one hand, improves *M. tuberculosis* clearance in already infected macrophages by promoting autophagy. On the other hand, it significantly suppresses both TNF-α and IFN-γ but leaves IL6 and IL10 unaffected, thereby interfering with further enhancement of anti-MTB activity (Ó Maoldomhnaigh et al., 2021). Moreover, lactate specifically upregulates matrix metalloproteinases that contribute to lung destruction in tuberculosis (Whittington et al., 2023), a process known to increase morbidity and mortality during TB treatment (Ong et al., 2014). The correlation between ES-Bifi and reduced red blood cell counts found in this study indirectly indicates decreased tissue oxygenation and, consequently, an increased risk of blood acidification in TB patients. Thus, the observed taxonomic and metabolic changes in the gut microbiome of TB patients may be a consequence of changes in host physiology — hypoxia in host tissues, a general increase in glycolysis, and lactate secretion during the immune response in TB (Llibre et al., 2021).

Thus, in this study, we first identified distinct patterns of bacterial communities, known as enterosignatures, in the gut microbiome of TB patients. We first showed that the gut microbiome of tuberculosis patients considerably differs from that of controls in terms of enrichment of ES-Bifi, ES-Esch, and ES-Firm, the abundance of which likely has a significant impact on host immunity beyond mycobacterial infection. Although enterosignatures are not discrete categories but are proposed as a method of stratification to reduce the complexity of microbial communities, they provide a valuable framework for understanding the relationship between gut microbiota and human health.

## Declarations

### Ethics approval and consent to participate

The studies involving humans were approved by the Ethics Committee of the Republican Clinical Antituberculous Dispensary (2019-0326). The studies were conducted in accordance with the local legislation and institutional requirements. The participants provided their written informed consent to participate in this study. Written informed consent was obtained from the individual(s) for the publication of any potentially identifiable images or data included in this article.

### Consent for publication

This article does not contain any individual person’s data in any form.

### Author Contributions

MY, and BY conceived and designed the study. LB, KK and IG were responsible for the selection of patients for the study, provided clinical services, extracted DNA from feces, and collected study data. LB and MY supervised the study. DS, AY, RA, and BY analyzed the data. BY, and RA were responsible for the methodology and provided statistical expertise. MY, LB, DS, and BY interpreted the results and drafted the manuscript. MY, DS, AY, and BY contributed to the writing of the manuscript. MY and BY took care of the editing of the manuscript and the final approval of the version for publication. All the authors have read and approved the final manuscript.

## Supporting information

https://docs.google.com/document/d/1j6moG2Y6mMKDuzGZs4TfEbQVxUWt3tGyqx4ZlbuHwrI/edit?tab=t.0

https://docs.google.com/spreadsheets/d/1LHnROUrOgnTYavZK_FQYhAD6LuVatMfvQF3uQ2Jt0Jc/edit?usp=drive_link

https://docs.google.com/spreadsheets/d/1ZqAiQW1aC9eqjv07XFwLygrlUnouhAOr/edit?usp=drive_link&ouid=102069768330273584533&rtpof=true&sd=true

## Acknowledgments

Not applicable

## Supplementary Material

The Supplementary Material for this article can be found online at:

1. Supplementary Materials
2. Supplementary Table 2
3. Supplementary Table 3

## Data Availability Statement

The shotgun metagenomic data used in this study are available in the NCBI database under BioProject accession codes PRJNA1366497 and PRJNA1061168. BioProject PRJNA1061168 stores shotgun metagenomic data for 70 samples collected and processed uniformly by our group but published as part of another case-control study (Yunusbaeva et al. 2024). Both data sets are available for research purposes upon request. Requests should be directed to the corresponding author, Bayazit Yunusbayev.

## Abbreviations

CLR: center log ratio
ES: enterosignature
FDR: False discovery rate
HIV: human immunodeficiency viruses
LDA: Linear Discriminant Analysis
LEfSe: Linear discriminant analysis effect size
MAGs: metagenomically assembled genomes
MTB: Mycobacterium tuberculosis
PCA: Principal coordinate analysis
TB: Tuberculosis

