## Supplementary material for "Bifidobacterium- and Escherichia-dominant ecological guilds shape distinct microbial metabolism in the gut microbiome of tuberculosis patients": https://docs.google.com/document/d/1j6moG2Y6mMKDuzGZs4TfEbQVxUWt3tGyqx4ZlbuHwrI/edit?tab=t.0

Supplementary Table 1. Clinical and demographic characteristics of the compared groups	2

Supplementary Figure 1. Power analysis of tuberculosis as a binary category	3

Supplementary Figure 2. Gut microbiome diversity and composition in ТB patients and healthy controls	4

Supplementary Figure 3. Taxonomic features of the gut microbiota of TB patients and controls	5

Supplementary Figure 4. Metabolic features of the gut microbiota of TB patients and controls	6

**Supplementary Table 1. Clinical and demographic characteristics of the compared groups**

|  | **TB patients (N =33)** | **Healthy control (N =47)** |
| --- | --- | --- |
| Female | 14/33 (42,4%) | 23/47 (49%) |
| Men | 19/33 (57,6%) | 25/47 (51%) |
| Age, years (Median (IQR) | 43 (34,5-55) | 40 (36-45) |
| Clinical forms of pulmonary tuberculosis | |  |
| Infiltrative TB | 19/33 (57,6%) | .. |
| Disseminated TB | 13/33 (39,4%) | .. |
| Caseous pneumonia | 1/33 (3,0%) | .. |

Data are median (IQR) (first and third quartiles) or n/N (%), unless stated otherwise. N ― number of individuals

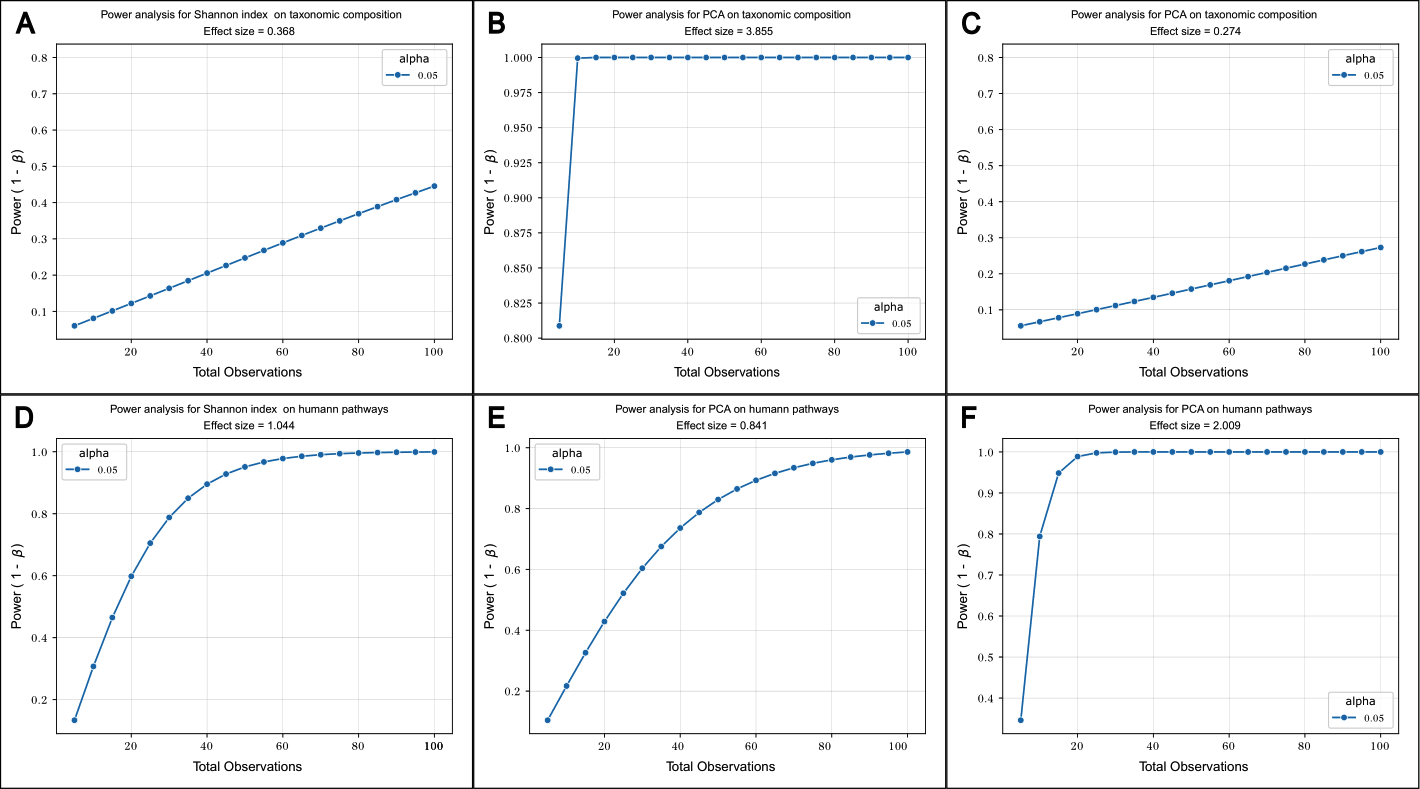

**Supplementary Figure 1. Power analysis of tuberculosis as a binary category.**

Power curves were computed using a significance level of 0.05.

The first (top) row shows: (A) Power estimates based on the effect size of tuberculosis (Cohen's d = 0.368) on taxonomic diversity (Shannon index); (B, C) Power estimates based on the effect of tuberculosis on taxonomic variation captured by PC1 (d = 3.855) and PC2 (d = 0.274).

The second (bottom) row shows: (D) Power estimates based on the effect size of tuberculosis (Cohen's d=1.044) on metabolic pathway diversity (Shannon index), as inferred using HUMAnN analysis; (E, F) Power estimates based on the effect size of tuberculosis on metabolic pathway variation captured by PC1 (d = 0.841) and PC2 (d = 2.009).

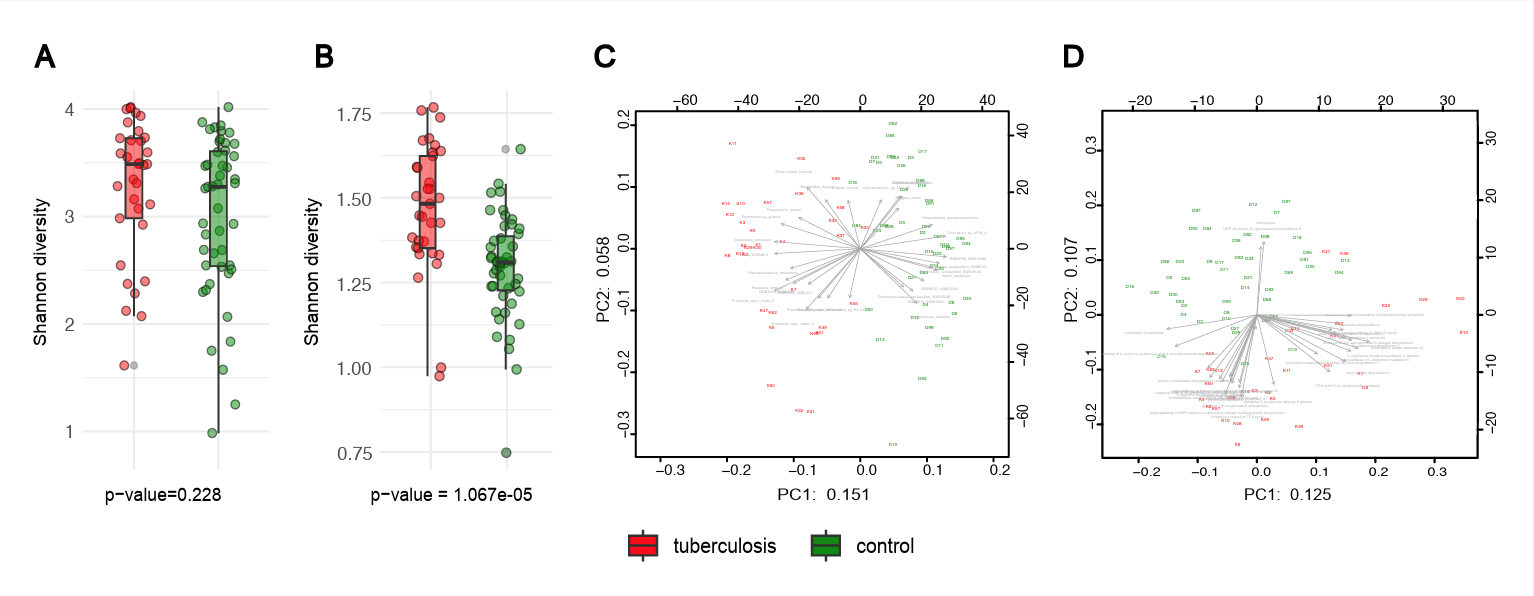

**Supplementary Figure 2. Gut microbiome diversity and composition in ТB patients and healthy controls.** The alpha diversity was assessed using the Shannon index. (A) A boxplot illustrating the Shannon diversity index, which represents the overall taxonomic diversity in the control and TB groups. No significant differences in diversity were observed between the TB and control groups (P > 0.05). (B) A boxplot illustrating the Shannon diversity index, which measures the diversity of metabolic pathways. The metabolic pathway diversity was significantly higher in TB patients than in the controls. The box represents the interquartile range; the horizontal line inside shows the median, the whiskers indicate the lower and upper quartiles, and points outside the whiskers indicate potential outliers. No significant differences in diversity between the TB and control groups (P > 0.05). (С, D) PCA analysis based on CLR-transformed relative abundances of bacterial taxa (C) and metabolic pathways (D). PC1 and PC2 are given with the proportion of total variance explained.

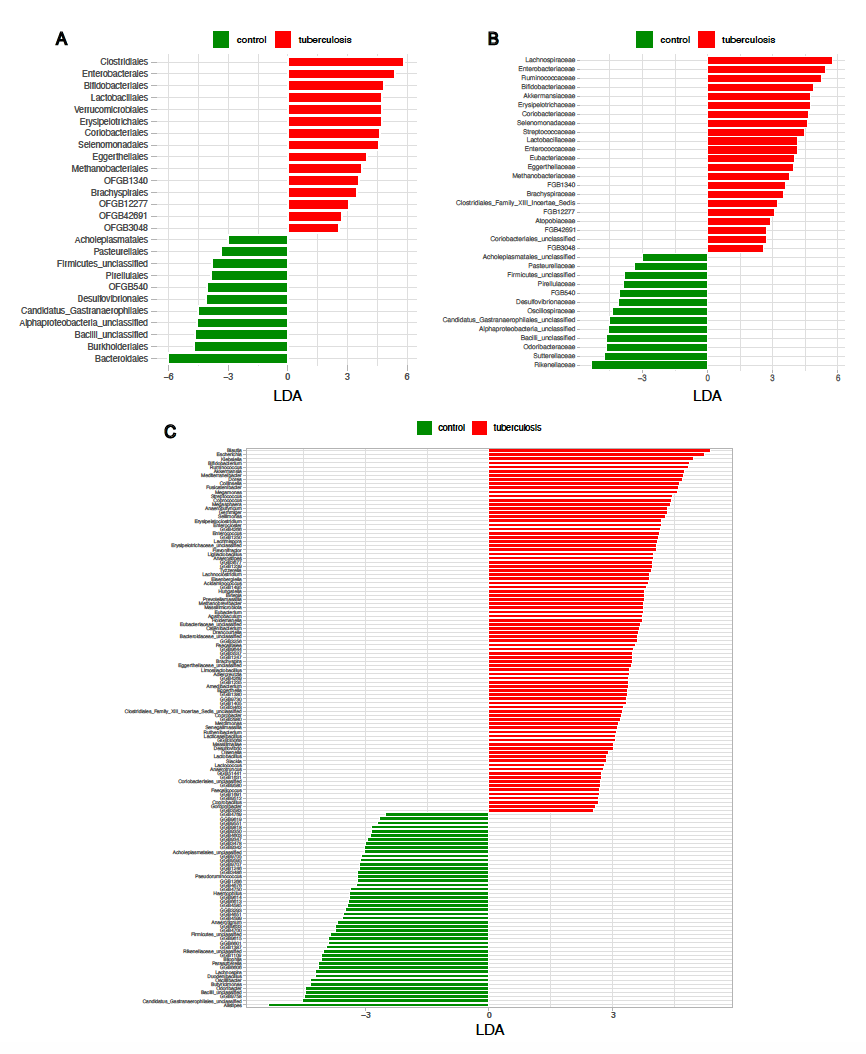

**Supplementary Figure 3. Taxonomic features of the gut microbiota of TB patients and controls.**

LDA score plots were generated using the LEfSe analysis. The length of the bar column represents the LDA score. Plots show the microbial taxa with significant differences between the TB patients (red) and healthy controls (green) at the order (A), family (B), and genus (C) level (LDA score > 2,5).

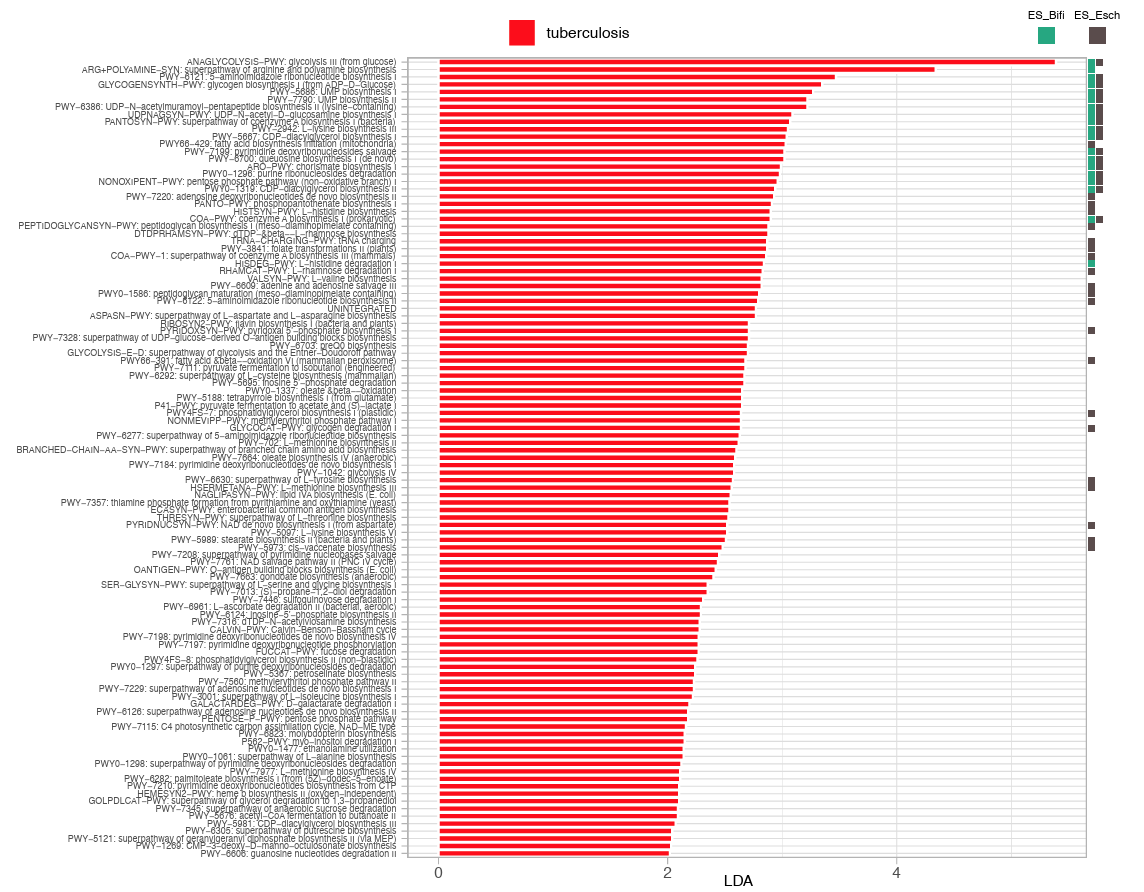

**Supplementary Figure 4.** **Metabolic features of the gut microbiota of TB patients and controls.**

LDA score plots were generated using the LEfSe analysis. The length of the bar column represents the LDA score. Plots show metabolic pathways with significant differences between the TB patients (red) (LDA score > 2). In the gut microbiota of TB patients, the top metabolic pathways demonstrating the greatest differentiation from healthy controls are predominantly associated with ES-Esch (represented by black squares) and ES-Bifi (indicated by turquoise squares).

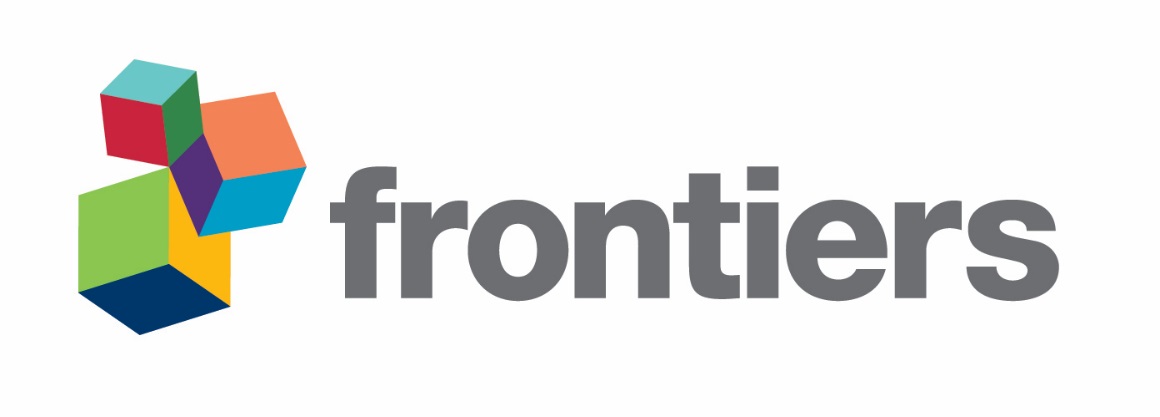
